# Cross-Dataset Transcriptomic Analysis Identifies Oxidative Stress–Inflammation Gene Networks Modulated by Nutrigenomic Interventions in Parkinson’s Disease

**DOI:** 10.64898/2026.05.05.723100

**Authors:** Masoumeh Rafiee, Faezeh Abaj, Reza Ghiasvand

## Abstract

Inflammation and oxidative stress (OS) are key to Parkinson’s disease (PD). We performed a cross-dataset integrative transcriptomic analysis to identify OS- and inflammation-related hub genes consistently dysregulated in PD and to explore gene–compound relationships using nutrigenomic studies using publicly available datasets. Four GEO datasets (GSE7621, GSE20141, GSE20146, GSE49036) were analysed to identify differentially expressed genes (DEGs), which were intersected with GeneCards OS–inflammation gene sets. Functional enrichment analyses, including gene ontology (GO), pathway over-representation analysis (ORA), and protein-protein interaction (PPI) analysis, were used to identify key pathways and hub genes. Gene–food bioactive compound (FBC) association was explored by integrating PD signatures with nutrigenomic profiles from NutriGenomeDB. We identified 183 DEGs in PD, enriched in synaptic, dopaminergic, OS, and inflammatory pathways. Intersection analysis yielded 26 OS-inflammation-related genes and 10 central regulators, including TH, DDC, SNCA, LRRK2, HSPB1, and HSPA1B. Integration with nutrigenomic datasets revealed opposing-direction transcriptional patterns, with several FBC-associated signatures showing lower expression of stress-related genes and higher expression of dopaminergic markers such as TH, GCH1, and DDC. Overall, this integrative analysis highlights OS–inflammation gene networks in PD and identifies candidate diet–gene associations that warrant further experimental and clinical validation.

## 1. Introduction

Parkinson’s disease (PD) is the second most common neurodegenerative disorder, characterised by motor symptoms such as resting tremor, muscle rigidity, bradykinesia, and postural instability [1]. The main pathological hallmark of PD is the degeneration of dopaminergic neurons in the substantia nigra (SN) and the accumulation of Lewy bodies in the surviving neurons [2]. However, the exact mechanisms underlying the pathogenesis of PD remain largely unknown, emphasising the need for extensive research into its causes, early diagnosis, and the development of effective therapeutic interventions.

Dopaminergic neuron degeneration is primarily caused by α-synuclein aggregation, impaired protein processing, oxidative stress (OS), proteasome dysfunction, mitochondrial damage, and neuroinflammation [3, 4]. OS and inflammation contribute to the pathogenesis of PD by promoting neurodegeneration, excitotoxicity, and axonal damage [5]. Various signalling pathways, including mitochondrial dysregulation, neuronal apoptosis, and neuronal inflammation, are affected by OS resulting from factors such as nuclear factor erythroid 2-related factor 2 (Nrf2) [6], nitric oxide synthase (NOS), manganese-dependent superoxide dismutase (MnSOD), cytochrome P450 (P450), HFE-related hemochromatosis, and methylenetetrahydrofolate reductase (MTHFR). Microglia, the primary innate immune cells in the central nervous system, become activated in response to inflammatory signals and OS, leading to the release of inflammatory cytokines [7].

Moreover, activated microglia produce reactive oxygen species (ROS), which contribute to neuronal inflammation and OS [8]. The interplay between chronic inflammation and OS exacerbates the progression of PD [9]. Notably, both genomic and environmental factors have a major impact on PD development, with diet among the most critical environmental determinants [10]. Several nutritional components, known as food bioactive compounds (FBCs), interact with gene expression and metabolic pathways, rendering them valuable for preventing and treating neurodegenerative diseases, particularly PD [11]. The most significant FBCs include carotenoids, organosulfur compounds, phenolic compounds, terpenoids, resveratrol, curcumin, fatty acids, saponins, probiotics, conjugated linoleic acid, and long-chain omega-3 polyunsaturated fatty acids [12].

While oxidative stress and inflammation are well-recognized contributors to PD, accumulating evidence shows that FBCs can enhance immune function and strengthen antioxidant defenses [13]. Nutrigenomics provides a mechanistic framework for understanding how dietary molecules modulate gene expression relevant to PD pathology. Essential micronutrients such as selenium and zinc activate selenoproteins and metallothionein-dependent enzymes (e.g., GPx, Cu/Zn-SOD), reducing oxidative DNA damage and supporting blood–brain barrier stability, whereas excess iron and copper promote ROS generation, α-synuclein aggregation, and neuronal apoptosis. Bioactive compounds (including resveratrol, curcumin, lipoic acid, and coenzyme Q10 (activate the Nrf2–ARE pathway, upregulating detoxification and mitochondrial biogenesis genes, while diet-derived factors also shape epigenetic regulation through DNA methylation and histone acetylation [14]. Collectively, these nutrigenomic mechanisms suggest that targeted dietary interventions may enhance neuroprotective gene expression, dampen inflammatory signaling, and potentially slow PD progression, supporting the broader concept of personalized nutrition for neurodegenerative disease management [15]. Cross-dataset transcriptomic integration has recently proven effective in identifying consistent gene-expression signatures across heterogeneous nutritional interventions, demonstrating the value of multi-study synthesis for uncovering convergent biological responses despite variability in study design and exposure conditions [16].

Although oxidative stress and inflammation are strongly implicated in Parkinson’s disease, the mechanistic roles of OS- and inflammation-related genes remain incompletely understood. To date, no comprehensive study has systematically identified key OS/inflammation-associated hub genes in PD or explored their associations with food bioactive compounds. This study aims to examine OS- and inflammation-related hub genes and their relationships with FBCs using an integrative bioinformatics approach. By combining transcriptomic and nutrigenomic data, this work provides a foundation for future mechanistic studies and precision nutrition research in PD.

## 2. Methods

### 2.1. Datasets and preprocessing

PD-related datasets were identified using the GEO database. The query included “Parkinson’s disease” and “neurodegenerative disease” with platform GPL570 (retrieved on 10/04/2025). These datasets include GSE7621, GSE20141, GSE20146, and GSE49036, encompassing studies that utilize microarray techniques alongside matched normal tissues. All datasets were restricted to the GPL570 (Affymetrix Human Genome U133 Plus 2.0) platform to ensure technical consistency across studies.

Specifically, GSE7621, GSE20141, and GSE49036 included SN samples from 16, 10, and 20 post-mortem PD patients, respectively, along with 9, 8, and 8 control samples. Meanwhile, GSE49036 featured 10 post-mortem PD globus pallidus internal samples and 10 samples from normal individuals. Although the SN and globus pallidus are anatomically distinct, both are integral components of the basal ganglia circuitry and are transcriptionally altered in PD.

Platform annotation files were obtained to assist in mapping probes to gene symbols. A list of oxidative stress and inflammation genes was sourced from the Gene Cards human gene database (https://www.genecards.org/) using the search terms “Oxidative Stress” and “Inflammation” (retrieved on 10/06/2025). Genes with a relevance score exceeding 7 were selected for further analysis. Finally, we obtained 1845 oxidative stress and 404 inflammation-related genes, of which 2019 genes were selected after removing common genes between the OS and inflammation gene sets for subsequent evaluation.

### 2.2. Differential expression genes analysis

Differential expression analysis was performed in R using the limma package (version 3.10.3). Raw CEL files from the GPL570 platform were background corrected and normalised using the Robust Multi-array Average (RMA) method. To minimise technical variability across datasets, batch effects were adjusted using the ComBat algorithm implemented in the sva package. After pre-processing, all samples were merged into a single expression matrix comprising 56 PD samples and 35 control samples. A design matrix was constructed modelling disease status (PD vs control) as the primary variable. Linear models were fitted using `lmFit`, followed by empirical Bayes moderation with `eBayes` to stabilize variance estimates across genes. Differential expression was assessed using moderated t-statistics, and p-values were adjusted for multiple testing using the Benjamini–Hochberg false discovery rate (FDR) method. Genes with |log2 fold change| ≥ 0.5 and FDR < 0.05 were considered significantly differentially expressed. Sample distributions before and after normalisation are shown in **Additional File 1.**

### 2.3. Functional enrichment analysis

#### 2.3.1 Gene ontology (GO) analysis and treemap visualisation

To summarise enriched GO categories into higher-level biological themes, semantic similarity between GO terms was computed using rrvgo (calculateSimMatrix, method = “Rel”). Redundant terms were collapsed into representative parent categories using reduceSimMatrix with a similarity threshold of 0.7. A treemap was generated to visualise major functional clusters, with tile size proportional to −log (q-value) and colours corresponding to parent GO categories, providing an intuitive overview of dominant biological processes enriched among the DEGs.

#### 2.3.2 Pathway enrichment

Pathway enrichment was performed using the human WikiPathways database for over-representation analysis (ORA). Significant genes were converted to Entrez IDs and tested for pathway enrichment. For enriched pathways of interest, contributing gene lists were extracted and mapped back to gene symbols and log fold-change values. To examine functional relationships among pathways, a connectivity network was constructed in Cytoscape using shared genes. Pathways were linked by edges when they contained overlapping DEGs, allowing identification of higher-order functional modules formed by gene sharing. Node colour reflected aggregated gene expression changes within each pathway based on log fold-change values.

#### 2.3.3 Gene-set enrichment using RummaGEO (GEO-derived signatures)

Complementary gene-set enrichment was performed using RummaGEO (https://rummageo.com/), a resource of transcriptional signatures derived from 29,294 GEO studies. Differentially expressed oxidative-stress and inflammation-related genes (DEOSIGs) were queried against the RummaGEO human gene-set library. Enriched signatures (adjusted p < 0.05) provided additional biological context from GEO-derived experimental contrasts.

#### 2.3.4 Protein-Protein Interaction (PPI) network analysis

The protein-protein interaction network analyzed through the DEGs and DEOSIGs dataset was constructed using the STRING database (https://string-db.org/) [17]. In the analysis of the PPI network, a confidence level of 0.4 or higher was set as the threshold. The PPI network data were visualized using Cytoscape software (version 3.9.1; https://cytoscape.org/) [18]. We utilized cytoHubba, a Cytoscape software plug-in, to extract the hub genes from the entire PPI network. Hub genes were identified using topological analysis in Cytoscape Network Analyzer. Nodes were ranked by degree centrality, and consistency was confirmed using betweenness and closeness centrality. Genes that ranked within the top quartile of degree and demonstrated high betweenness centrality were defined as hub genes.

### 2.4. Impact of FBCs on the expression of genes related to PD

Based on bioinformatics analysis, we extracted a supplementary file containing FBCs and their GSE IDs from the NutriGenomeDB databases associated with cell lines. NutriGenomeDB consist of manually curated gene sets defining nutrigenomics experiments available in the GEO database (http://www.nutrigenomedb.org/). The Gene Expression Browser module was used, based on a curated set of differentially expressed genes obtained from 287 nutrigenomics experiments (retrieved on 15/11/2021). All experiments are linked to their original GEO sources. After careful screening, 67 appropriate studies with GSE IDs were selected (listed in **Additional File 2**).

#### 2.4.1 Gene–FBC interaction heatmaps

To evaluate whether FBCs modulate PD-related genes in the opposite direction, two heatmaps were generated: one for genes upregulated in PD but downregulated following FBC treatment, and one for genes downregulated in PD but upregulated after FBC exposure. For each gene–compound pair, the PD and FBC log fold-change values were visualised side by side using a red–blue diverging scale to reflect the direction and magnitude of regulation. FBC tiles included the compound name and corresponding expression change to enable direct comparison with the PD profile.

## 3. Results

### 3.1. Identification of differentially expressed genes (DEGs)

Using limma-based linear modelling with FDR adjustment and |log2FC| ≥ 0.5, we identified 183 differentially expressed genes (48 upregulated and 135 downregulated) in PD compared to controls (**Additional File 3**). The volcanic plot of the 183 DEGs is presented in **Fig. 1**.

**Fig. 1:**
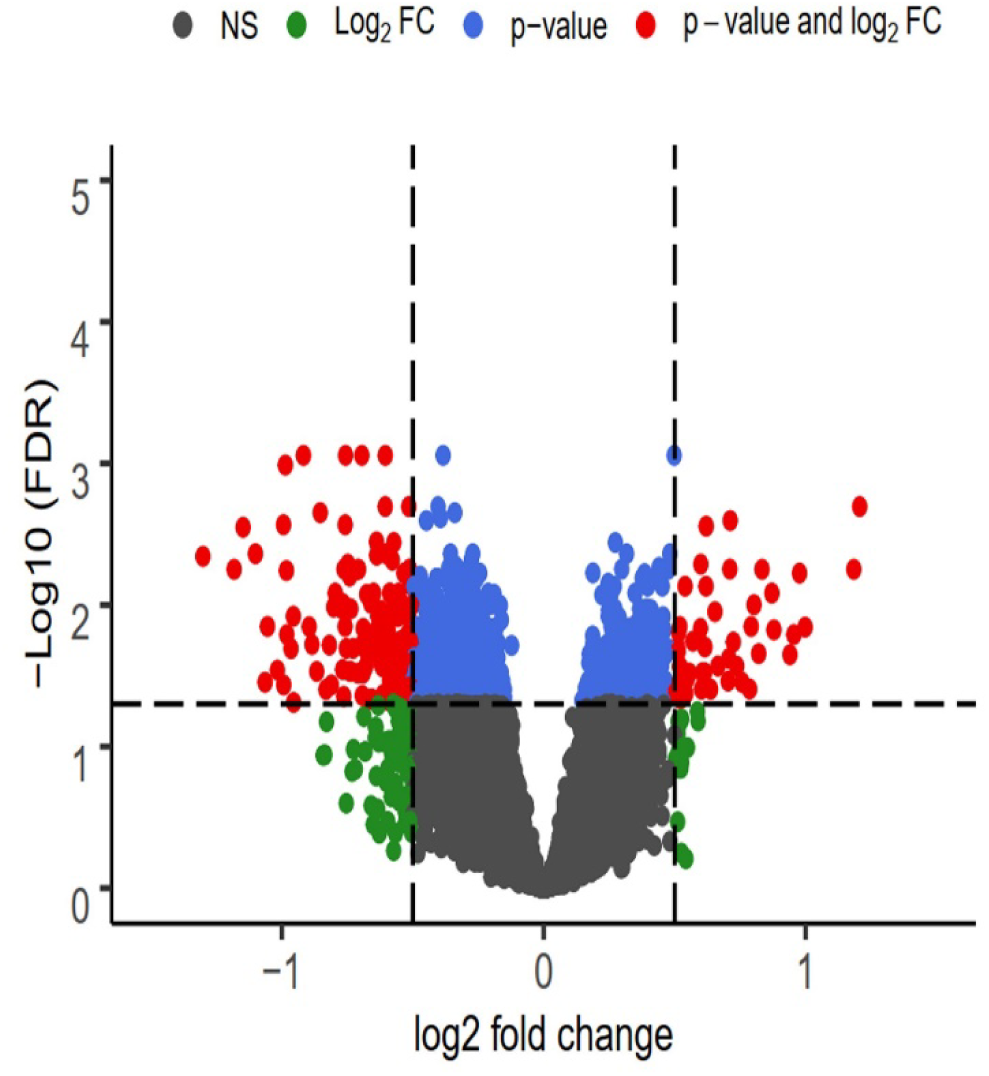
Volcano plot showing 183 differentially expressed genes (DEGs) identified across the merged PD datasets.

### 3.2. Treemap visualisation of GO term clusters

Semantic similarity reduction using identified several high-level functional themes. The dominant clusters included synaptic signalling, vesicle trafficking, autophagy regulation, response to metal ions, oxidative stress, axonogenesis, learning and cognition, and response to temperature or reactive oxygen species. Overall, the treemap showed that PD DEGs are enriched in biological processes regulating synaptic structure, neurotransmission, vesicle transport, neuronal development, and stress response mechanisms (**Fig.2**).

**Fig 2:**
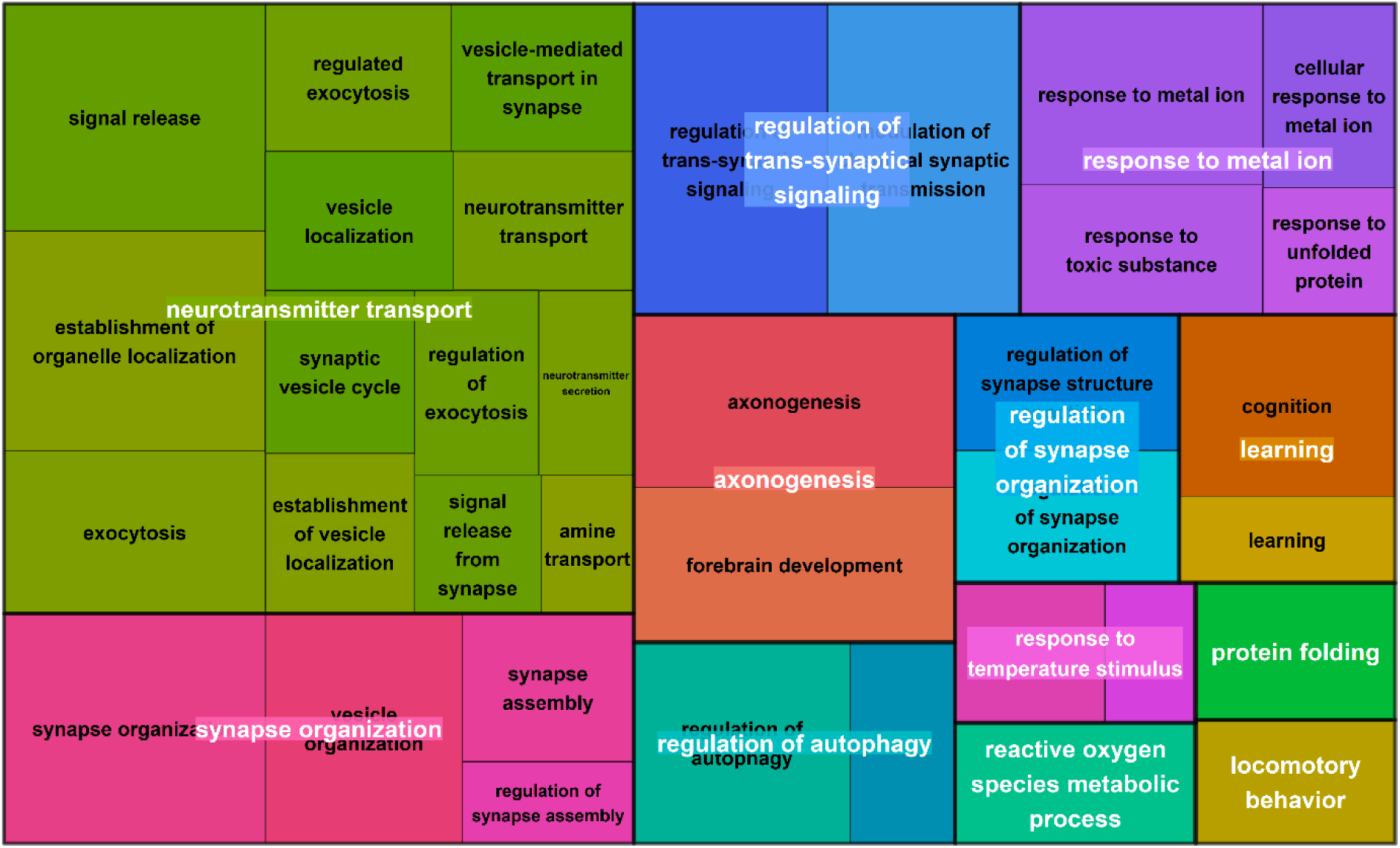
Treemap visualization of GO term clusters derived from the 183 DEGs, summarizing high-level functional themes.

### 3.3 Pathway enrichment analysis of DEGs in PD samples

ORA detected several significantly enriched WikiPathways terms among the PD-related DEGs. Detailed information on all pathways is provided in **additional file 4**. The most prominent enrichment was in Dopaminergic neurogenesis (WP2855; adjusted p-value <0.001, q = 5.55 × 10), driven by eight key genes: SLC18A2, ALDH1A1, EN1, RET, DDC, TH, SLC6A3, and NR4A2. Additionally, the pathway Neurotransmitter disorders (WP4220; adjusted p-value = 5.55× 10; q = 5.26 × 10) showed significant enrichment, with four overlapping genes: SLC18A2, DDC, TH, and SLC6A3.

The Parkinson’s disease pathway (WP2371; adjusted p-value = 8.86 × 10 ³, q = 7.71 × 10 ³) was identified as a significant module, enriched with genes such as SNCA, DDC, TH, SLC6A3, and LRRK2. Additionally, there was evidence of broader neurodevelopmental involvement suggested by the enrichment in the ADHD and autism /ASD pathway (WP5420; adjusted p-value = 1.11× 10^02^, q = 9.96 × 10 ³), which includes 14 genes such as AGTR1, AKR1C3, SLC25A32, PRKAR2B, KYAT3, CNTN4, DDC, TH, and SLC6A3.

Metabolic dysregulation was indicated by the enrichment of Amino acid metabolism (WP3925; adjusted p-value = 4.24 × 10 ², q = 3.80 × 10 ²), involving GSS, ALDH1A1, BCAT1, DDC, TH, and AUH. Additionally, oxidative-stress–related cell death was demonstrated by Ferroptosis (WP4313; adjusted p-value = 4.97 × 10 ², q = 4.46 × 10 ²), supported by HMOX1, HSPB1, AKR1C3, KIT, and GCH1.

Several genes appeared across multiple pathways, functioning as hubs. For example, TH and DDC were common to dopaminergic neurogenesis, neurotransmitter disorders, and PD pathways; ALDH1A1 was involved in both dopaminergic neurogenesis and amino acid metabolism; and AKR1C3 connected neurodevelopmental signalling, metabolism, and ferroptosis. These shared genes highlight the convergence of dopaminergic dysfunction, synaptic impairment, metabolic disturbance, and oxidative stress within the PD transcriptomic landscape (**Fig.3**).

**Fig 3:**
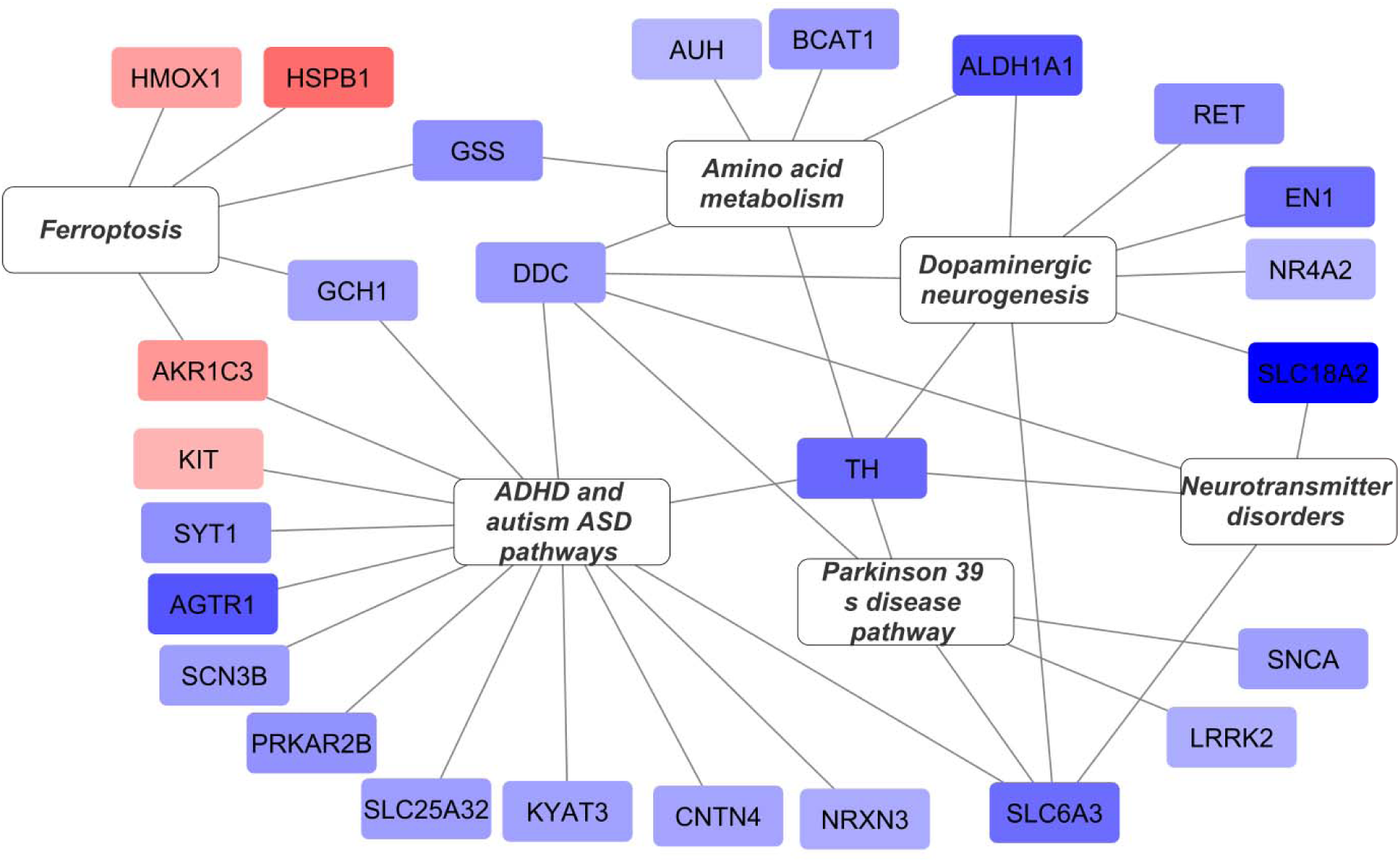
Over-representation analysis (ORA) of PD-associated DEGs

### 3.4. Oxidative stress and inflammation-related DEGs (DEOSIGs)

Venn diagram analyses revealed the overlap between DEGs and gene lists related to oxidative stress and inflammation (**Fig. 4A**). After comparing 183 DEGs with 2019 genes associated with oxidative stress and inflammation, we identified 26 differentially expressed oxidative stress– and inflammation-related genes (DEOSIGs). Fourteen genes were downregulated: SNCA (log2FC = −0.656, adj.P = 0.0161), TH (−1.017, 0.0289), GCH1 (−0.614, 0.0430), GSS (−0.757, 0.0009), AGTR1 (−1.148, 0.0028), TTC19 (−0.506, 0.0277), GBE1 (−0.896, 0.0142), LRRK2 (−0.553, 0.0384), SLC6A3 (−0.993, 0.0367), ALDH1A1 (−1.182, 0.0056), DDC (−0.680, 0.0273), VRK1 (−0.578, 0.0048), CCK (−0.532, 0.0428), and SCN3B (−0.657, 0.0461).

Twelve genes were upregulated: HMOX1 (0.654, 0.0112), HSPB1 (0.998, 0.0144), STIP1 (0.706, 0.0239), CRYAB (0.614, 0.0301), BAG3 (0.879, 0.0150), HSPA1B (1.206, 0.0020), AKR1C3 (0.711, 0.0056), SERPINH1 (0.821, 0.0221), NUPR1 (0.803, 0.0100), CXCR4 (0.871, 0.0082), HSPA6 (1.184, 0.0056), and BMI1 (0.502, 0.0421).

#### 3.4.1 PPI network analysis of DEOSIGs

To explore protein–protein interactions, three PPI networks were constructed. First, separate networks were generated for the 48 upregulated DEGs (**Fig. 4B**) and 135 downregulated DEGs (**Fig. 4C**). Similarly, a PPI network was constructed with the 26 DEOSIGs (12 up-regulated and 14 down-regulated genes), which included 26 nodes and 50 edges (**Fig. 5D**). This DEOSIG-specific network highlighted key interaction modules enriched for oxidative stress and inflammation-related regulators.

**Fig. 5.**
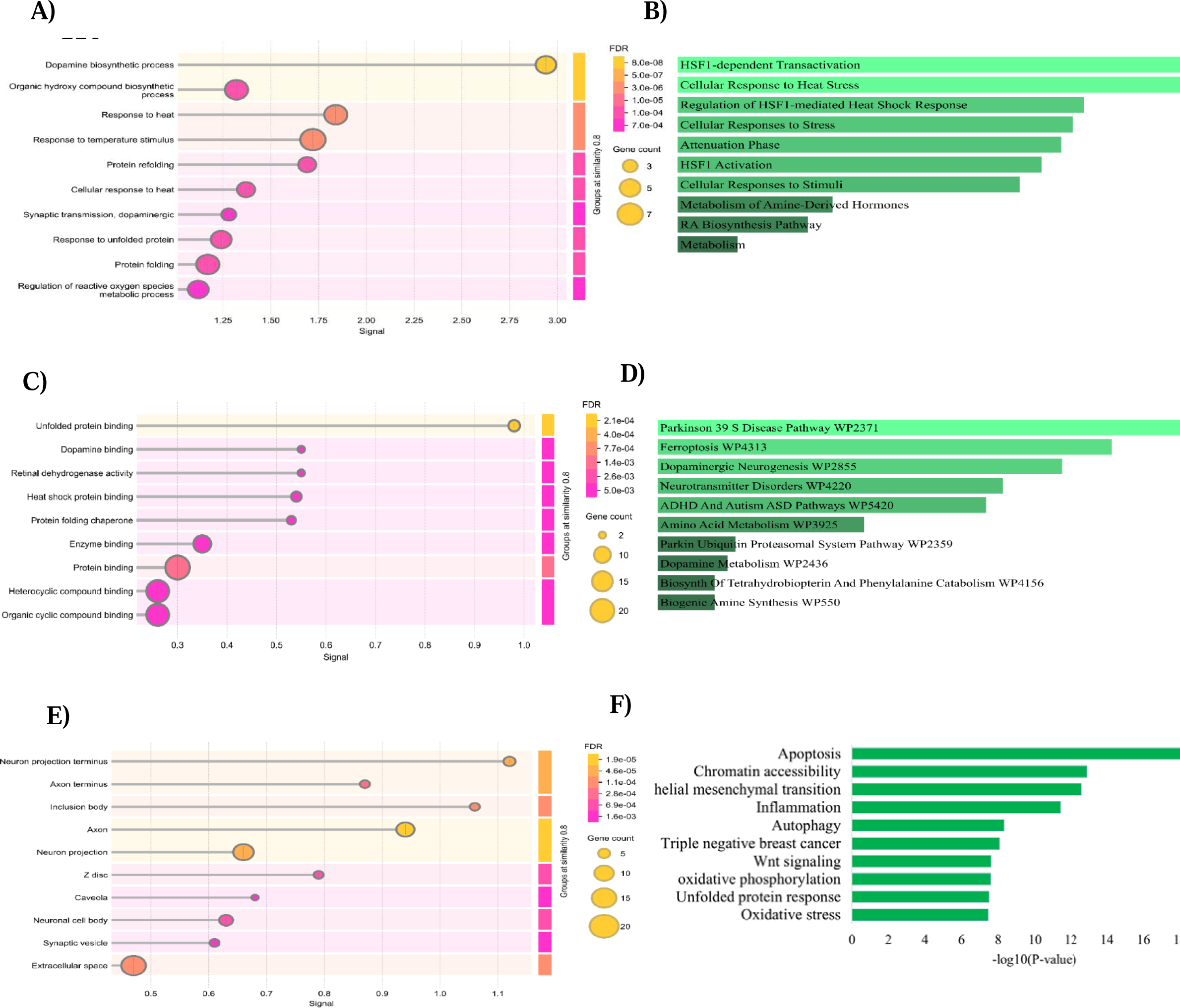
Functional annotation of the 26 overlapping genes (DEOSIGs). **(A)** Biological processe (BP). **(B)** Cell composition (CC). **(C)** Molecular function (MF). **(D)** Reactome Pathways 2024. **(E)** WikiPathways 2024 Human**. (F)** RummaGEO.

#### 3.4.2 Functional enrichment of DEOSIGs

For a more detailed investigation, we elucidated these 26 DEOSIGs in GO, Reactome pathways and WikiPathways 2024 human. In GO BP, DEOSIGs were mainly enriched in dopamine biosynthetic process, response to heat, response to temperature stimulus, protein refolding, cellular response to heat, organic hydroxy compound biosynthetic process, synaptic transmission, dopaminergic, response to unfolded protein, protein folding and regulation of reactive oxygen species metabolic process (**Fig. 5A**). For GO CC, DEOSIGs are primarily enriched in neuron projection terminus, axon terminus, inclusion body, axon, neuron projection, Z disc, caveola, neuronal cell body, synaptic vesicle and extracellular space (**Fig. 5B**). GO MF term was mostly enriched in unfolded protein binding, dopamine binding, retinal dehydrogenase activity, heat shock protein binding, protein folding chaperone, enzyme binding, protein binding, heterocyclic compound binding and organic cyclic compound binding (**Fig. 5C**). Reactome pathways commentary revealed that the DEOSIGs were mainly enrolled in several pathways, including HSF1-dependent transactivation, cellular response to heat stress, regulation of HSF1-mediated heat shock response, cellular responses to stress, attenuation phase, HSF1 activation, cellular responses to stimuli, metabolism of amine-derived hormones, RA biosynthesis and metabolism (**Fig. 5D**). The result of WikiPathways analysis showed that DEOSIGs were primarily involved in Parkinson 39 S disease, ferroptosis, dopaminergic neurogenesis, neurotransmitter disorders, ADHD and Autism ASD, amino acid metabolism, parkin ubiquitin proteasomal system, dopamine metabolism, biosynth of tetrahydrobiopterin and phenylalanine catabolism and biogenic amine synthesis (**Fig. 5E**). The results of RummaGEO showed that the top 10 biological processes and pathways contain apoptosis, chromatin accessibility, epithelial-mesenchymal transition, inflammation, autophagy, triple-negative breast cancer, wnt signalling, oxidative phosphorylation, unfolded protein response, oxidative stress and immune response were highly correlated with DEOSIGs (**Fig. 5F**).

### 3.5 Identification of hub genes

We obtained the top 10 hub downregulated-DEGs (TH, SLC18A2, DDC, SLC6A3, SNCA, EN, KCNJ6, GCH, LRRK2 and CCK), upregulated-DEGs (DNAJB1, HSPA1B, HSPB1, HSPA6, HSPA1L, BAG3, STIP1, PTGES3 and AHSA1) and DEOSIGs (HSPB1, HSPA1B, SNCA, TH, DDC, HSPA6, BAG3, SLC6A3, LRRK2 and STIP1) according to their degree score (**Fig. 6A-C**).

**Fig. 6.**
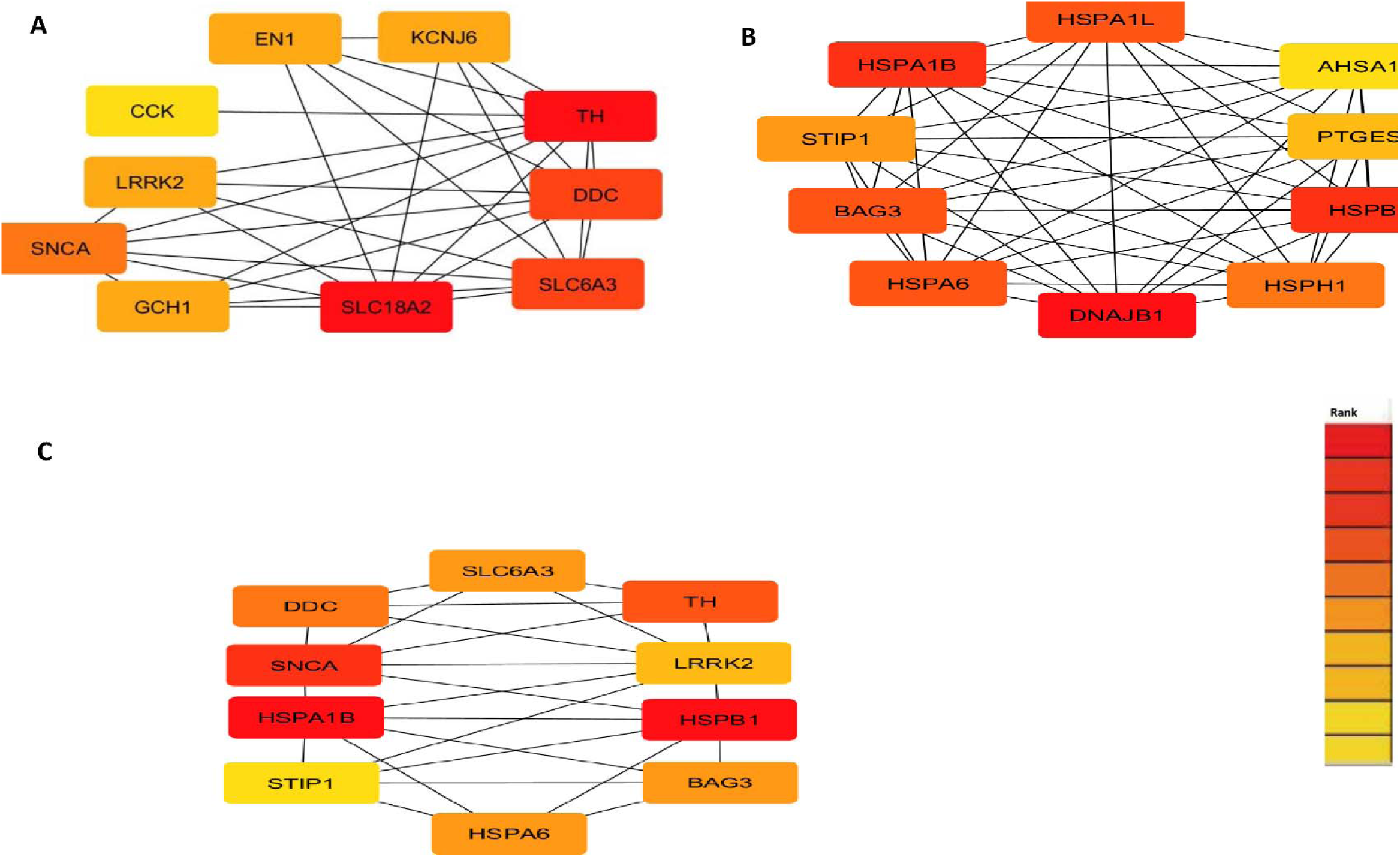
The top 10 hub genes screened by degree values. **(A)** Down-regulated DEGs **(B)** up-regulated DEGs **(C)** DEOSIGs. Different colors indicate the rank of degree.

### 3.6 Opposing interaction between FBCs and PD-related genes

To determine whether FBCs counteract transcriptional alterations observed in PD, we compared PD-related differential expression patterns with gene expression responses to 67 nutrigenomic interventions. Two major categories of opposing interactions were identified:

#### 3.6.1 Genes upregulated in PD but downregulated after FBC treatment

A total of 19 genes showed increased expression in PD but were consistently decreased after exposure to specific FBCs, as illustrated in the heatmap (**Fig 7A**).

**Fig. 7:**
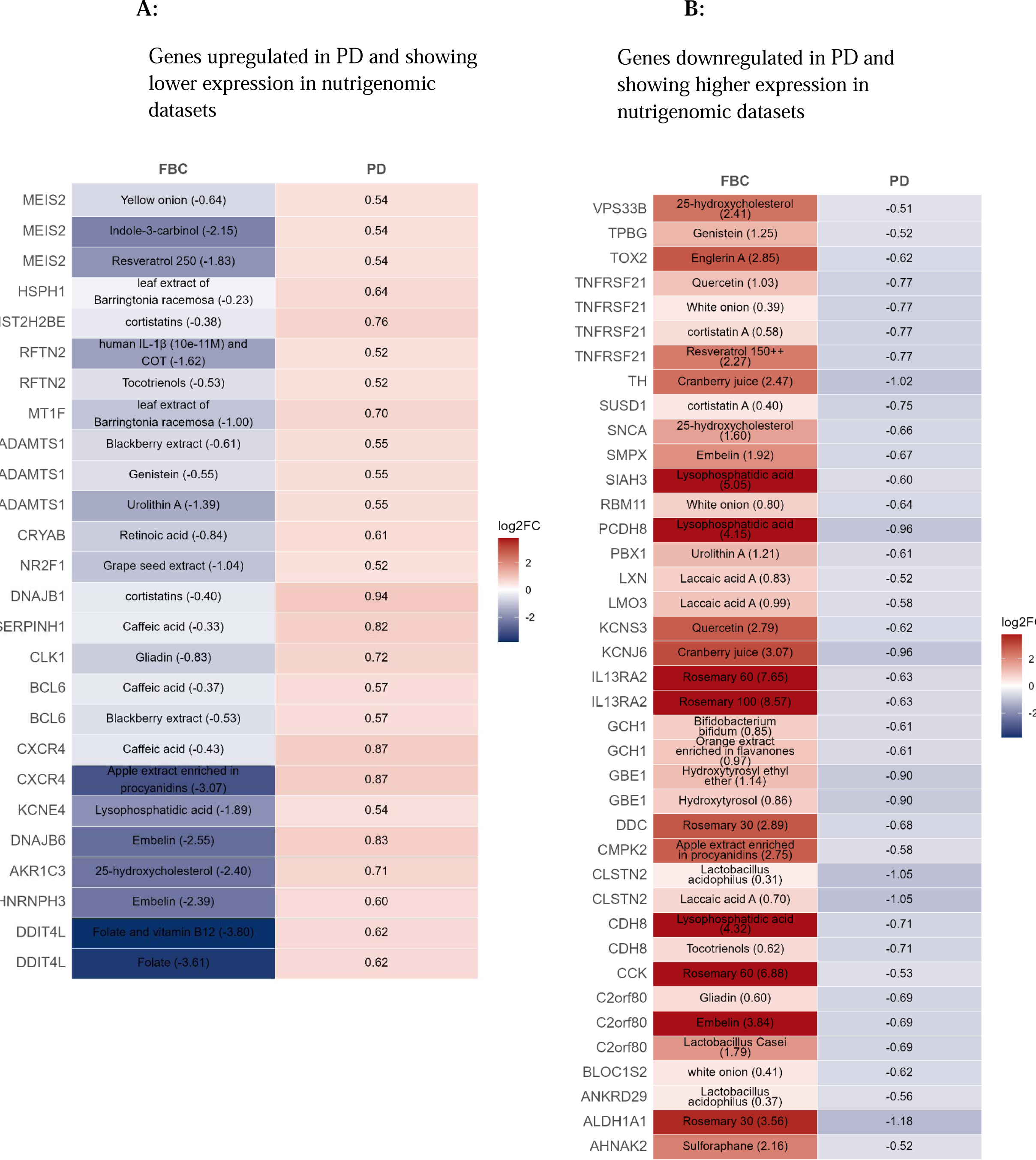
Opposing-direction transcriptional patterns between PD-associated signatures and nutrigenomic datasets

DDIT4L, which was elevated in PD, was downregulated by both Folate and Folate plus Vitamin B12. Several stress-response genes, including HNRNPH3 and DNAJB6, were suppressed by Embelin, while AKR1C3 responded to 25-hydroxycholesterol. The potassium channel gene KCNE4 showed decreased expression following lysophosphatidic acid treatment. Inflammation- and chemotaxis-related genes, such as CXCR4, exhibited reduced expression following apple extract enriched in procyanidins and caffeic acid, while CXCR5 was also downregulated by caffeic acid. The transcriptional regulator BCL6 was decreased by blackberry extract and caffeic acid.

Metabolic genes, including CLK1 and SERPINH1, were downregulated by gliadin and caffeic acid, respectively. Chaperone-associated genes, including DNAJB1 and HIST2H2BE, responded strongly to cortistatins, whereas NR2F1 was reduced by grape seed extract. Oxidative stress–related CRYAB levels declined after retinoic acid treatment. Meanwhile, ADAMTS1 expression was lowered by urolithin A, genistein, and blackberry extract. Detoxification genes MT1F and the adaptor protein RFTN2 also decreased following treatments with Barringtonia racemosa leaf extract, tocotrienols, and human IL-1β combined with COT. Additionally, the heat-shock regulator HSPH1 was reduced after Barringtonia racemosa leaf extract exposure. The neurodevelopmental gene MEIS2 was downregulated by resveratrol 250 and by indole-3-carbinol combined with yellow onion.

Together, these genes are involved in PD-related stress, chaperone, inflammatory, and metabolic pathways, which seem to be actively inhibited by various bioactive compounds.

#### 3.6.2 Genes downregulated in PD but upregulated after FBC treatment

Conversely, 29 genes showed reduced expression in PD but increased following FBC intervention and are shown in a heatmap (**Fig. 7 B**).

Several genes exhibiting lower expression in PD tissue, including KCNJ6, TH, GCH1, and DDC, showed increased expression in selected nutrigenomic datasets. Specifically, KCNJ6 was associated with cranberry juice, TH with rosemary extract, GCH1 with orange flavanone extract, and DDC with Bifidobacterium bifidum PRL2010. Similarly, immune-related genes such as IL13RA2 and TNFRSF21 displayed higher expression in response to certain compounds, including genistein, urolithin A, and blackberry extract. Additional neuronal and metabolic genes, including CLSTN2, ANKRD29, C2orf80, AHNAK2, GBE1, SMPX, LMO3, LXN, CDH8, SIAH3, and PCDH8, also exhibited opposing-direction transcriptional patterns relative to PD-associated signatures. Neurodevelopmental regulators (PBX1, TPBG, RBM11, BLOC1S2, and TOX2) showed similar trends. Although the specific associations varied across compounds and experimental systems, these observations suggest that certain food bioactive compounds may influence transcriptional networks relevant to dopaminergic, immune, metabolic, and neuronal processes. However, these findings should be interpreted as hypothesis-generating associations rather than evidence of functional restoration or therapeutic efficacy.

## 4. Discussion

PD is the most prevalent motor disorder affecting the brain [19]. Several studies indicate that OS and neuroinflammation play crucial roles in PD development. However, the precise mechanisms by which OS- and inflammation-related genes contribute to the pathophysiology of PD remain unclear. Through an integrative transcriptomic analysis combining multi-study DEGs, OS–inflammation gene sets, and nutrient-response datasets, we identified molecular networks associated with dopaminergic dysfunction, metabolic imbalance, and stress-response pathways, and transcriptional patterns in which several FBCs exhibited opposing regulation relative to PD-associated alterations.

The GO and pathway enrichment analyses revealed that Parkinson’s disease involves not only the loss of dopaminergic neurons but also extensive disturbances across synaptic, metabolic, oxidative, and inflammatory pathways. The notable enrichment of processes related to synapses—such as synapse structure, vesicle cycling, neurotransmitter transport, and exocytosis—suggests widespread alterations in presynaptic and postsynaptic signalling. This supports the known degeneration of nigrostriatal dopaminergic neurons and points to broader synaptic impairments affecting multiple neural systems.

Metabolic dysfunction was particularly prominent, especially in amino acid metabolism and catecholamine synthesis. The repeated enrichment of TH, DDC, ALDH1A1, and GCH1 across dopaminergic neurogenesis, neurotransmitter imbalances, and amino acid metabolic pathways highlights the close relationship between dopaminergic pathways, amino acid metabolism, and cellular energy processes in PD.

Alongside synaptic and metabolic changes, we observed a marked increase in oxidative and inflammatory activities. Clusters related to ROS metabolism, heat-shock response, protein refolding, metal-ion response, autophagy, and proteostasis highlight known disease mechanisms like mitochondrial dysfunction, defective protein quality control, and neuroinflammation. Additionally, ferroptosis-related gene signatures were enriched in our analysis, suggesting a potential role for iron-dependent oxidative pathways in PD, which warrants further experimental validation. Beyond traditional PD biology, pathways related to neurodevelopment, such as axonogenesis, learning, cognition, and forebrain development, showed significant enrichment. The presence of transcriptional modules linked to ADHD and ASD suggests a wider vulnerability of neural connection systems or shared molecular features across neurodevelopmental and neurodegenerative disorders [20, 21].

### 4.1 Intersection of ORA pathways and DEOSIGs identifies key molecular convergence points

Thirteen genes were notably shared between the ORA-enriched pathways and the 26 DEOSIGs, suggesting a potential overlap between oxidative stress, inflammatory, and PD-associated molecular processes. These overlapping genes (SNCA, TH, GCH1, GSS, AGTR1, LRRK2, SLC6A3, ALDH1A1, DDC, SCN3B, HMOX1, HSPB1, and AKR1C3) represent candidate molecular links connecting dopaminergic dysfunction, metabolic dysregulation, oxidative stress, and inflammatory signalling. A multitude of these genes are pivotal to dopamine biosynthesis (TH, DDC, ALDH1A1, GCH1) [22], vesicular transport and synaptic homeostasis (SLC6A3, SCN3B, SNCA) [23], oxidative stress response and detoxification (GSS, HMOX1, HSPB1, AKR1C3) [24], and neuroinflammatory pathways (AGTR1, LRRK2) [25]. Their co-enrichment across synaptic, metabolic, and stress-response pathways highlights coordinated involvement of metabolic, redox, and inflammatory processes in PD pathology. SLC6A3 (Solute Carrier Family 6 Member 3) is a sodium-dependent dopamine transporter gene [26]. The overexpression of SLC6A3 in dopamine (DA) neurons leads to dopamine accumulation in presynaptic neurons, resulting in dopamine peroxidation and neurotoxicity due to increased free radical production [27]. In our analysis, SLC6A3 expression was reduced in PD samples. Given that SLC6A3 is predominantly expressed in dopaminergic neurons, this reduction may partly reflect dopaminergic neuron loss rather than transcriptional suppression within surviving neurons [28].

Leucine-rich repeat serine/threonine-protein kinase 2 (LRRK2) is a multifunctional kinase involved in autophagy, stress responses, and immune regulation [29]. Pathogenic LRRK2 mutations are typically linked to increased kinase activity, promoting microglial pro-inflammatory cytokine production and dopaminergic vulnerability [30], whereas pharmacological inhibition reduces inflammatory signalling in experimental models [31]. In contrast, our analysis detected reduced LRRK2 mRNA expression in PD substantia nigra samples. Given that transcript levels do not necessarily reflect kinase activity and may be influenced by neuronal loss, treatment effects, or disease stage, the biological significance of this reduction remains uncertain and should be interpreted cautiously.

Furthermore, we identified 5 hub upregulated-DEGs (HSPB1, HSPA1B, HSPA6, BAG3, and STIP1) and 5 hub downregulated-DEGs (TH, DDC, SLC6A3, SNCA, and LRRK2) that showed strong associations with OS and inflammation. GO enrichment analysis revealed that DEOSIGs were mainly involved in cellular responses to heat and stress, HSF1-mediated heat shock regulation, dopamine biosynthesis, protein folding and refolding, oxidative stress regulation, and dopaminergic synaptic transmission. These findings align with the established roles of OS and inflammation in PD pathogenesis. Our findings suggest that the identified genes are enriched in pathways previously implicated in PD, including neuroinflammation, oxidative stress, and mitochondrial dysfunction [32–34]. ROS production in PD mainly results from dopamine metabolism, mitochondrial complex I deficiency, and chronic neuroinflammation [35–38]. Altered expression of these genes has been associated with immune signalling dysregulation, chronic inflammation, and blood–brain barrier dysfunction in previous studies [39]. Converging evidence suggests that dopamine deficiency stems from the death of dopaminergic neurons in the substantia nigra pars compacta (SNpc), attributed to the activation of inflammatory pathways and increased OS [40].

### 4.2 Biological relevance of Hub genes in parkinson’s disease

The identified hub genes show varying levels of mechanistic and associative relevance to PD. Strong mechanistic evidence from previous studies supports SNCA, LRRK2, TH, and DDC, which are directly involved in dopamine biosynthesis, neuronal survival, and α-synuclein aggregation core processes in PD pathology. Similarly, HSPB1 and HSPA1B have been reported to exert neuroprotective effects by preventing misfolded protein accumulation and enhancing aggregate clearance. Moderate functional evidence supports BAG3 and STIP1, both implicated in protein quality control, BAG3 through selective autophagy and STIP1 via HSP70/HSP90 chaperone coordination where they mitigate α-synuclein toxicity in model systems, though direct human evidence remains limited. SLC6A3 (dopamine transporter) shows reduced expression following dopaminergic degeneration and may contribute to PD susceptibility through altered dopamine reuptake, while HSPA6 appears indirectly related through general stress-response pathways. Collectively, these findings indicate that hub genes such as SNCA, LRRK2, TH, DDC, HSPB1, and HSPA1B have established mechanistic roles, whereas BAG3, STIP1, SLC6A3, and HSPA6 are functionally or associatively linked candidates. Integrative molecular network approaches that combine transcriptomic profiling with protein–protein interaction analysis have been successfully applied in other complex disorders to identify novel candidate genes and tissue-specific dysregulation patterns [41]. However, hub status reflects network connectivity and prioritization within the interaction network rather than confirmed biological importance. Therefore, although these genes occupy central network positions, their functional relevance to PD pathogenesis requires further experimental validation. Distinguishing these levels of evidence refines the biological interpretation of PD-related gene networks and highlights targets for future mechanistic investigation.

### 4.4 Interaction between hub genes and food bioactive compounds

An important aspect of this study was the integration of nutrigenomic data to explore associations between FBC-related transcriptional signatures and PD-associated gene expression patterns. By comparing PD transcriptional signatures with responses to 67 nutrigenomic interventions, we observed two major classes of opposing-direction transcriptional patterns: genes upregulated in PD but suppressed by FBCs, and genes downregulated in PD but exhibiting higher expression in selected nutrigenomic datasets. Many PD-elevated genes belonged to stress-response, chaperone, inflammatory, and metabolic pathways, and lower expression of these genes in FBC-associated signatures reflected an opposing transcriptional direction relative to PD-associated signatures [42, 43]. Folate-based treatments decreased DDIT4L expression; Embelin and 25-hydroxycholesterol downregulated HNRNPH3, DNAJB6, and AKR1C3; and anti-inflammatory polyphenols, including caffeic acid and blackberry extract, suppressed CXCR4/CXCR5 and BCL6 [44, 45]. Additional compounds such as urolithin A, retinoic acid, tocotrienols, and Barringtonia racemosa extract reduced expression of stress-related and metabolic genes, suggesting potential modulation of stress-related and metabolic pathways associated with PD.

Conversely, several genes that exhibited lower expression in PD tissue including key dopaminergic markers TH, KCNJ6 reflecting its polyphenol-mediated antioxidant potentials [46], GCH1 and DDC exhibited increased expression in selected nutrigenomic datasets by specific FBCs such as cranberry juice, rosemary extract, orange-flavanone extract, and Bifidobacterium bifidum. Probiotics have been reported in experimental systems to influence inflammatory and oxidative pathways relevant to dopaminergic neuron vulnerability [73].

Tyrosine hydroxylase (TH) is the primary rate-limiting enzyme involved in DA synthesis within neuronal cells. The reduction of TH in PD is bilateral. Firstly, the expression of the TH gene is diminished under conditions related to OS, particularly in DA neurons located in the substantia nigra [47]. Secondly, the degeneration and loss of TH-containing dopamine neurons can lead to TH deficiency in the brain [48]. Cellular and molecular studies have reported that the downregulation of the TH gene may exacerbate oxidative stress and neuroinflammation in PD [49]. Accordingly, current evidence has shown that substances with antioxidant and anti-inflammatory properties may contribute to an increase in the expression of TH in models PD [50].

DOPA-decarboxylase (DDC) is a pyridoxal phosphate-dependent enzyme that converts L-DOPA to DA. Based on experimental observations, the depletion of dopamine in neural cells from PD samples enhanced the expression of the DDC gene [51]. However, our transcriptomic analysis identified lower DDC expression in PD samples, possibly due to pharmacological treatment or enzyme dysfunction [52–55]. Consistent with our study, experimental evidence suggests that DDC acts as a cellular antioxidant and potential therapeutic target in PD [56].

GCH1, a key enzyme in tetrahydrobiopterin (BH4) synthesis, functions as an antioxidant and modulator of ferroptosis defense [57]. It also serves as a cofactor for tyrosine hydroxylase, essential for dopamine biosynthesis, and its reduced expression may contribute to PD pathogenesis [58]. Experimental studies suggest that flavonoid-rich interventions can enhance BH4 synthesis and modulate oxidative stress and neuroinflammatory pathways [59], supporting a potential role for GCH1 regulation in redox balance. Notably, clinical evidence also suggests that higher dietary purine intake, via uric acid–mediated antioxidant effects, is associated with improved motor outcomes after deep brain stimulation in PD, further supporting the relevance of redox modulation in disease progression [60].

Together, these findings indicate that diverse FBCs are associated with opposing-direction transcriptional patterns relative to PD-associated gene signatures across dopaminergic, inflammatory, metabolic, and neurodevelopmental pathways. While these observations do not demonstrate functional restoration or neuroprotection, they suggest that certain bioactive compounds may modulate gene networks implicated in PD pathophysiology. Further experimental studies are required to determine whether these transcriptional associations translate into meaningful effects on neuronal survival or function.

However, it is important to note that antioxidant and dietary supplementation trials in Parkinson’s disease have yielded inconclusive clinical outcomes [63], with several studies failing to demonstrate clear disease-modifying effects [64, 65]. These discrepancies highlight the complexity of translating molecular antioxidant mechanisms into clinical benefit and underscore the need for cautious interpretation of nutrigenomic findings.

Although this integrative transcriptomic analysis provides mechanistic insight into oxidative stress– and inflammation-associated gene networks in PD, the findings remain exploratory. Furthermore, the oxidative stress and inflammation gene sets were derived from GeneCards, and therefore the identified DEOSIGs should be interpreted as candidate genes associated with these processes rather than definitive mechanistic regulators.

Heterogeneity among included studies, limited clinical metadata, and the absence of experimental validation, such as RNA quality, as in post-mortem studies, warrant cautious interpretation. Because substantia nigra and globus pallidus tissues were analysed together to identify convergent PD-associated signatures, region-specific transcriptional differences cannot be excluded. Therefore, the identified DEGs and pathways should be interpreted as shared alterations across affected basal ganglia regions rather than region-restricted changes. Furthermore, nutrigenomic datasets were derived from heterogeneous experimental systems encompassing diverse cell types, compounds, treatment doses, and exposure durations. This biological and methodological heterogeneity limits direct comparison with PD brain tissue and complicates interpretation of gene–compound relationships. Therefore, the observed opposing-direction transcriptional patterns should be considered hypothesis-generating associations rather than validated mechanistic reversal or evidence of therapeutic efficacy. Future studies incorporating region-specific analyses, perturbation-based validation, longitudinal multi-omics [66], and in vivo models will be required to determine functional and translational relevance of these findings.

## 5. Conclusion

This study identified crucial hub genes linked to oxidative stress and inflammation that form the molecular basis of Parkinson’s disease, demonstrating that several genes are regulated by FBCs in food. Cross-dataset analysis revealed consistent perturbations in dopaminergic, metabolic, synaptic, and stress-response pathways, with nutrigenomic interventions exhibiting transcriptional patterns opposite to those associated with PD. These findings highlight diet-gene interactions as a framework for future mechanistic investigation.

## Abbreviations

PD: Parkinson’s disease;
FBCs: food bioactive compounds;
SN: substantia nigra;
DEGs: Differentially expressed genes;
DEOSIGs: Differentially expressed oxidative stress and inflammation genes;
OS: Oxidative stress;
ORA: Over Representation Analysis;
GEO: Gene Expression Omnibus;
GO: Gene Ontology;
KEGG: Kyoto Encyclopedia of Genes and Genomes;
PPI: Protein-protein interaction;
STRING: the Search Tool for the Retrieval of Interacting Genes;
FDR: false discovery rate.

## Acknowledgements

This study is the result of a research project No. 3400392 approved by the Vice-Chancellor for Research of the Isfahan University of Medical Sciences.

## Author contributions

Masoumeh Rafiee: Designed the research, Writing - Original draft preparation. Faezeh Abaj: Writing –Reviewing and Editing, Visualization and Validation. Reza Ghiasvand: designed the research, Supervision, Project administration.

## Data availability

Datasets are available through the first author upon reasonable request.

## Declaration of interest

The authors declare no conflicts of interest.

---

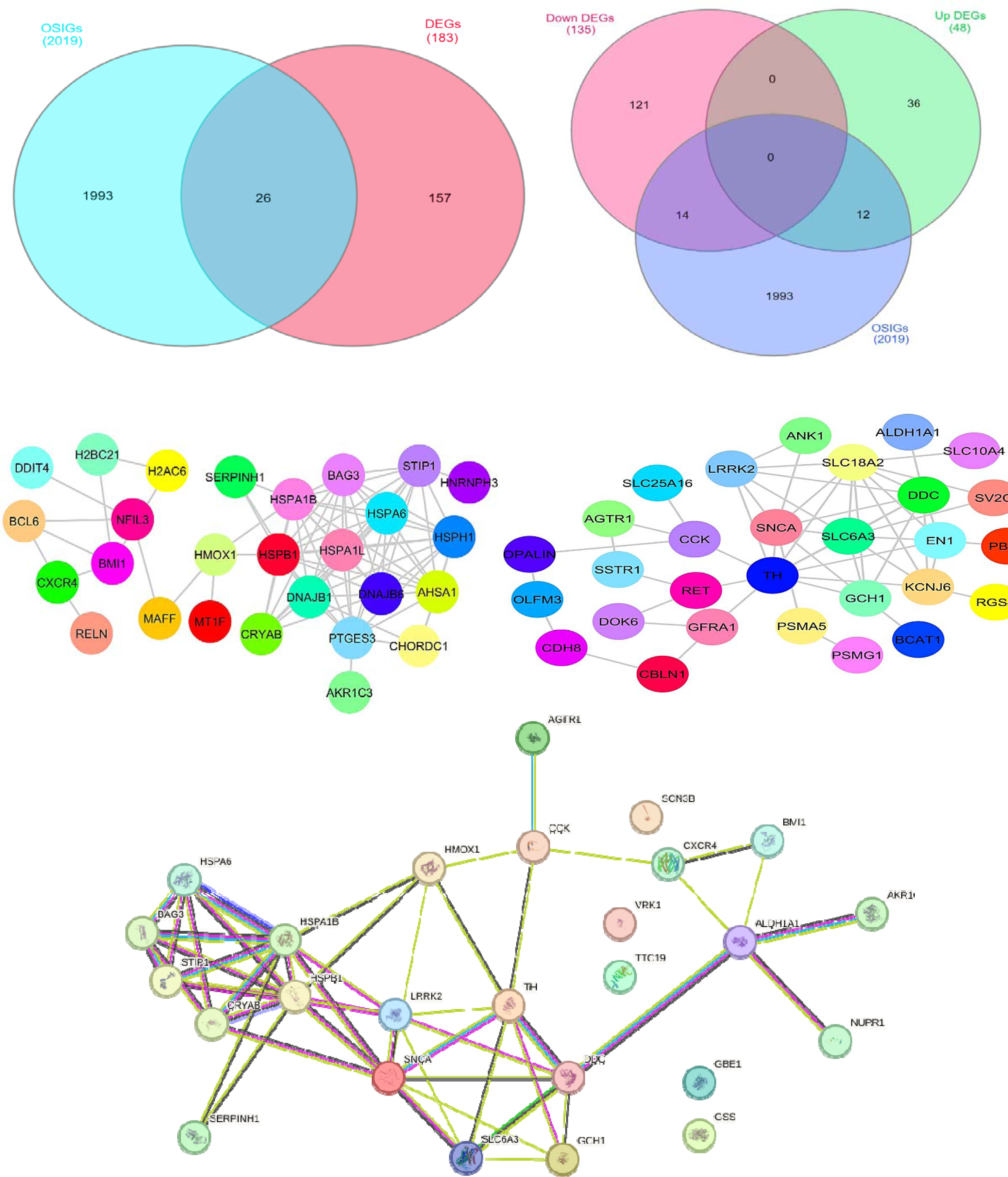

## Notes

### Competing Interest Statement

The authors have declared no competing interest.

### Summary of Updates

Major revision following peer-review feedback. The manuscript text, Abstract, Discussion, Conclusion, and figure legends were revised to improve clarity, reduce overinterpretation, and expand discussion of study limitations. Author list is also updated.

