## Supplementary figures and images for "Cross-Dataset Transcriptomic Analysis Identifies Oxidative Stress–Inflammation Gene Networks Modulated by Nutrigenomic Interventions in Parkinson’s Disease"

### Additional file 1.pdf

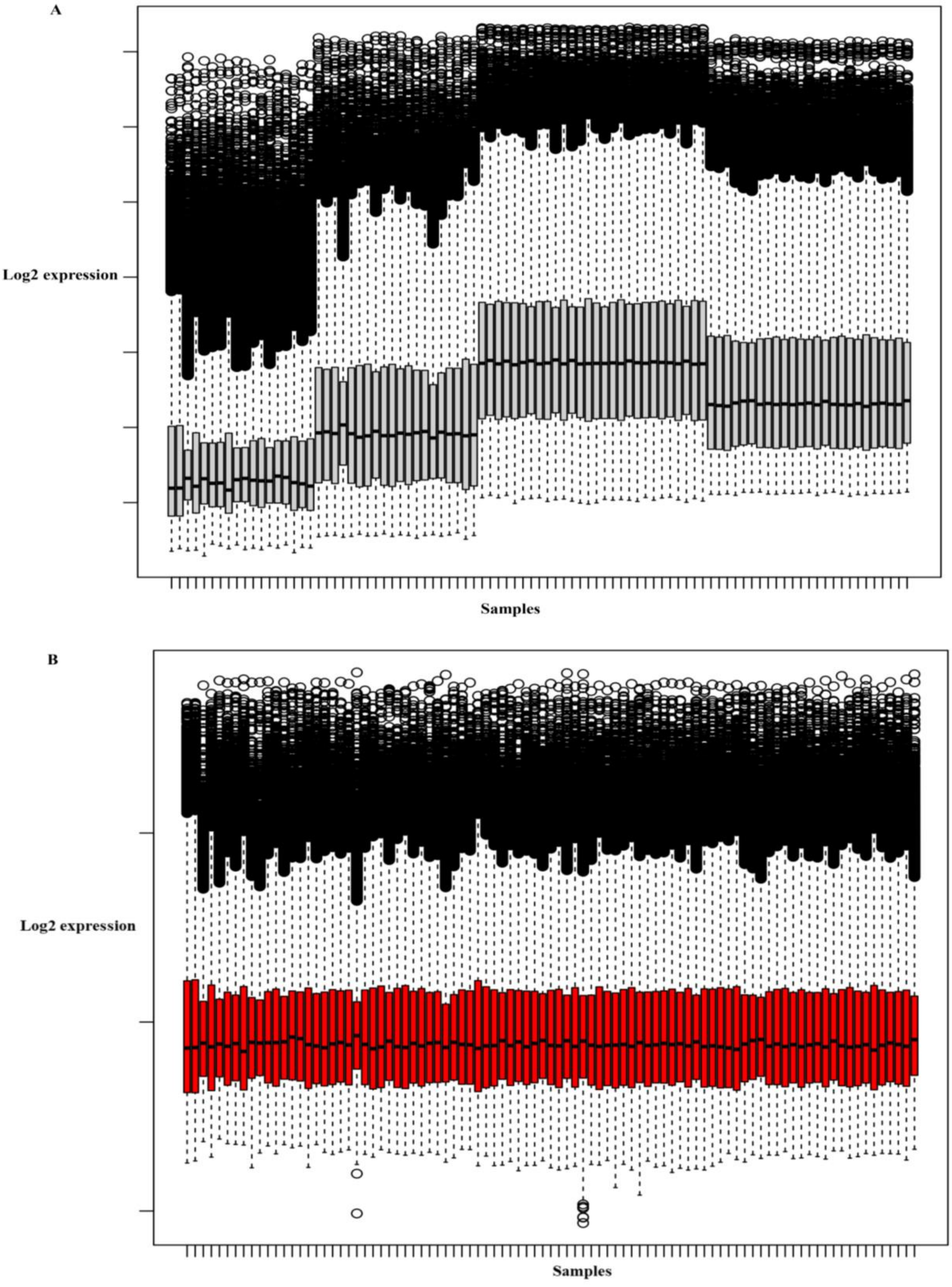


Data distribution before (**A**) and after (**B**) batch effect correction
